# BacteReason: A Reasoning Model for Antimicrobial Resistance Prediction

**DOI:** 10.64898/2026.06.04.730229

**Authors:** Yuna Oikawa, Shuichi Kawashima, Akira R. Kinjo, Yosuke Demizu, Ryo Tamura, Koji Tsuda

**Affiliations:** Graduate School of Frontier Sciences, The University of Tokyo, Kashiwa, Chiba 277-8561, Japan; Database Division for Life Science, BioData Science Initiative, National Institute of Genetics, Kashiwa, Chiba 277-0871, Japan; Anima Machina G.K., Kita-ku, Osaka 530-0001, Japan; Division of Organic Chemistry, National Institute of Health Sciences, Kawasaki, Kanagawa 210-9501, Japan; Center for Basic Research on Materials, National Institute for Materials Science, Tsukuba, Ibaraki 305-0044, Japan; RIKEN Center for Advanced Intelligence Project, Chuo-ku, Tokyo 103-0027, Japan

## Abstract

The rapid global spread of antimicrobial resistance (AMR) has placed unprecedented pressure on clinical decision-making. Machine learning predictors of antibiotic susceptibility exist, but their lack of mechanistic grounding limits credibility. We present BacteReason, a reasoning large language model (LLM) that predicts bacterial susceptibility to a target antibiotic, together with a mechanistic rationale. BacteReason is obtained by fine-tuning an open-weight LLM on clinical susceptibility data augmented with rationales that explain the molecular mechanisms. These rationales are produced by a proprietary teacher LLM prompted to explain known susceptibility outcomes. The teacher is interfaced via TogoMCP with a collection of biomedical knowledge-graph databases, grounding each reasoning step in retrieved evidence. On an extrapolation benchmark, BacteReason achieves a relative improvement of 43% over the untuned baseline and 38% over the same base LLM fine-tuned without rationales, demonstrating that reasoning supervision improves prediction accuracy.

## Introduction

Antimicrobial resistance (AMR) was attributed to 1.27 million deaths in 2019 [1]. One of its causes is inappropriate antibiotic stewardship [2]. Although antimicrobial susceptibility testing is recommended in clinical practice, obtaining results requires several days, a delay that can be critical for patient outcomes [3]. Antibiotics are therefore often prescribed empirically before culture-based results return, forcing clinicians to balance adequate patient coverage against preserving long-term antibiotic efficacy. Inappropriate empirical choices accelerate the selection and spread of resistant strains, further compounding the AMR crisis. Resolving this tension requires personalised susceptibility predictors.

Machine learning approaches to AMR prediction have shown promise [4, 5], but such predictions are not grounded in molecular mechanism. This is a fundamental limitation: resistance and susceptibility arise from cascades of molecular interactions, including enzyme inactivation, target site modification, and efflux pump expression. When a model reasons through these mechanisms, users would place more trust in the predictions. This motivates the use of large language models (LLMs), which can perform multi-step reasoning [6]. LLMs have been shown to provide rationales in biological domains such as medical question answering [7], disease diagnosis [8], and genomics [9], and here we apply the same idea to AMR prediction. There are at least two challenges to address. The first is financial cost. Reasoning with proprietary large models is much more expensive than simple question answering [10], and may not be affordable at the bedside, particularly in resource-limited settings where AMR is most pressing. The second is the incorporation of expert biological knowledge. Even the largest language models to date lack detailed knowledge of bacterial biology, including genomes, metabolism, and cellular processes. Without access to such knowledge, the mechanistic rationale provided by an LLM can lack detail or factual accuracy.

We present BacteReason, a reasoning LLM that predicts susceptibility to a target antibiotic, given a biosample’s antibiogram against other antibiotics (Figure 1). It also generates a rationale for its prediction, namely, the inferred molecular mechanism. To keep the running cost small, BacteReason is obtained by fine-tuning a small, open-weight LLM (Figure 2). The training dataset is generated from 1,875 clinical susceptibility records systematically derived from the AMR Portal database [11]. Each record is first transformed into a question–answer pair such that the question asks whether a biosample is susceptible to an antibiotic, and the answer is “susceptible” or “resistant”. A proprietary “teacher” LLM then uses chain-of-thought reasoning to augment each question–answer pair into a question– rationale–answer triplet, where the rationale explains the inferred molecular mechanism by which the answer follows from the question.

**Figure 1:**
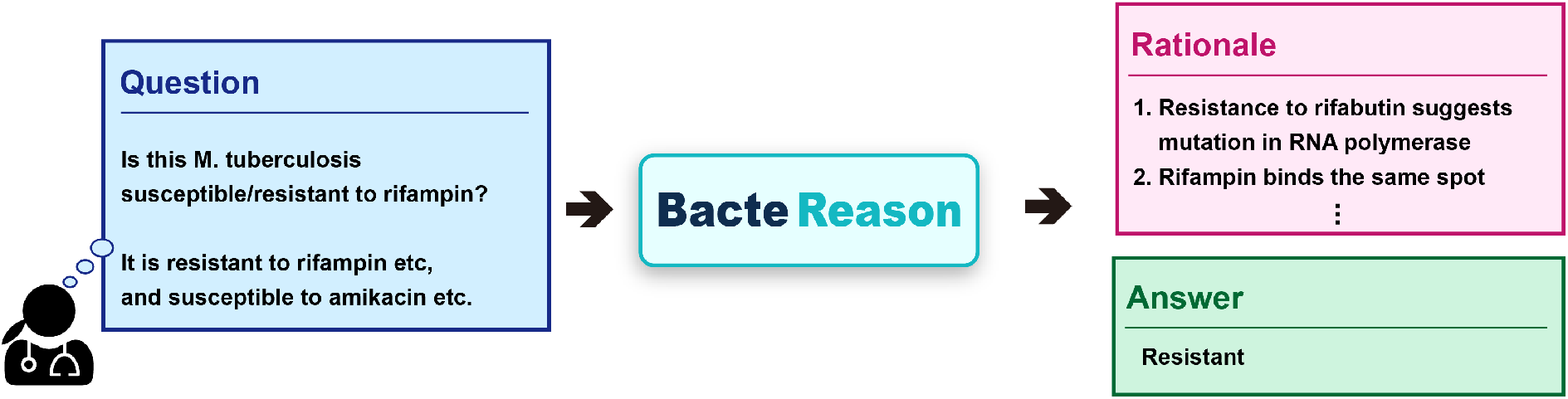
BacteReason at inference. Given a question comprising a clinical biosample’s antibiogram and the name of the target antibiotic, BacteReason outputs a chain-of-thought rationale followed by a binary susceptibility prediction.

**Figure 2:**
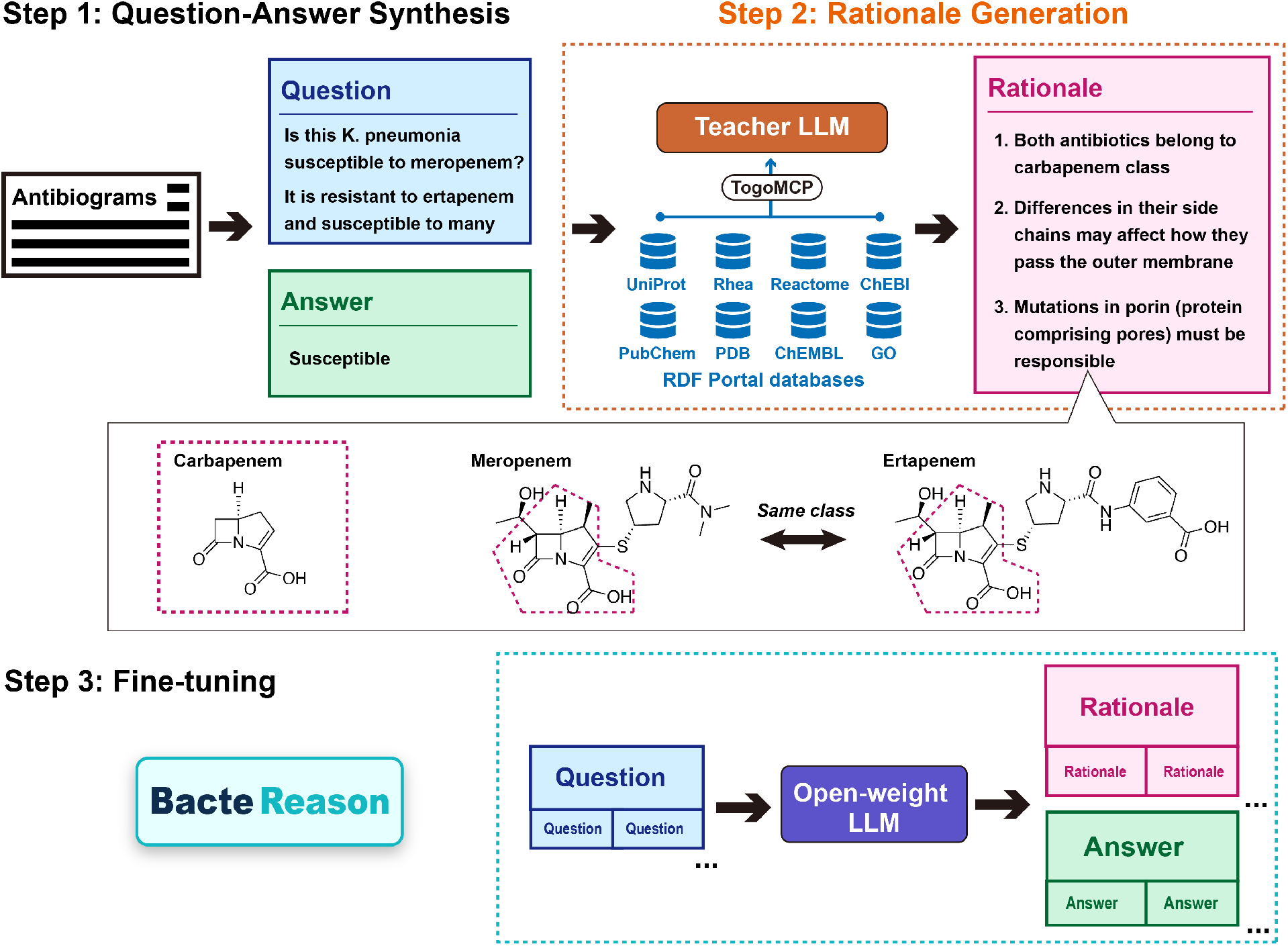
Training BacteReason. Step 1 (Question–Answer Synthesis): each clinical susceptibility record from the AMR Portal database is reformatted into a question–answer pair, where the question presents the biosample’s antibiogram with one antibiotic held out together with the name of that antibiotic, and the answer is the ground-truth susceptibility label. Step 2 (Rationale Generation): each pair is passed to a proprietary teacher LLM that, through TogoMCP, can autonomously query the RDF Portal biomedical knowledge graphs to construct a molecular rationale linking the antibiogram to the answer. Step 3 (Fine-tuning): the resulting question–rationale–answer triplets are used to fine-tune an open-weight LLM, producing BacteReason.

An unaugmented teacher LLM generates a rationale based only on internal knowledge. To create a high-quality rationale, the LLM should be able to query a variety of biomedical databases to collect necessary information proactively. We employed TogoMCP [12], a Model Context Protocol server to interface the teacher LLM with a collection of biomedical databases including UniProt, Rhea, Reactome, PubChem, ChEBI, ChEMBL, PDB, and GO. TogoMCP accesses knowledge-graph versions of 23 databases through eight SPARQL endpoints. It makes each database LLM-accessible by supplementing it with *semantic* and *syntactic* contexts. The former is used to inform the LLM about the content of the database. For example, ChEMBL is described as follows: *Manually curated database of bioactive molecules with drug-like properties containing* ^∼^*1*.*9M compounds, 1*.*9M assays, 21M+ bioactivity measurements, and comprehensive drug development data. Major entities include small molecules, proteins, targets, assays, activities, documents, drug mechanisms, and drug indications. Enables queries for compound–target–activity relationships, mechanism of action, drug repositioning, and pharmaceutical research*. The latter instructs the LLM on how to write the code to query the database. Through TogoMCP, the LLM can select appropriate databases and issue relevant queries.

On a benchmark of 200 clinical isolates, BacteReason achieved an error rate of 18% in susceptibility prediction, a relative improvement of 43% over the untuned baseline. When TogoMCP is removed, prediction accuracy drops sharply, showing that access to biological knowledge is essential for rationale quality. When the rationales are removed from the training data altogether, the accuracy deteriorates further. These findings show that grounding agentic LLMs in biological knowledge bases improves their reasoning, and the approach should generalise to other biological prediction tasks.

## Results

### Training BacteReason

BacteReason’s reasoning ability is acquired in three steps: question–answer pair synthesis, rationale generation, and fine-tuning (Figure 2). In Step 1, 1,875 question–answer pairs are systematically synthesized from randomly selected clinical biosample antibiograms (together with MIC values and genotypes where available) on the AMR Portal database [11].

In Step 2, each pair is reformatted as an instruction prompting the teacher LLM, Claude Opus 4.5 (extended thinking) [13], to generate a rationale linking the question to its answer (Supplementary Text S1), yielding a question–rationale–answer triplet. During this rationale generation, the teacher LLM is granted access to biomedical knowledge graphs through TogoMCP [12]. Through this single interface, the teacher can autonomously retrieve information from chemistry, structural biology, genetics, and microbiology as needed when writing a rationale.

In Step 3, the 1,875 triplets are used to fine-tune QwQ-32B [14], an open-weight LLM, with the loss computed on the rationale and answer tokens while question tokens are masked (see Methods for fine-tuning details). The fine-tuned LLM, supervised to produce a rationale that leads to the correct answer, is called BacteReason.

We next highlight two teacher-generated rationales that illustrate counterintuitive susceptibility outcomes. The first addresses biosample SAMEA104075919, a *Klebsiella pneumoniae* isolate susceptible to meropenem despite the resistance to ertapenem, a drug of the same carbapenem class. While no single paper accounts for this specific biosample, retrieved structural data from ChEBI suggested that ertapenem carries a bulkier side chain than meropenem, which may contribute to ertapenem’s greater sensitivity to porin constriction. This prompted the hypothesis that porin modifications — such as *ompK36* mutations or reduced *ompK35* /*ompK36* expression, as identified by the PDB database — could differentially restrict carbapenem entry. Doumith et al. [15] lend support to this, having demonstrated that restoring functional porins in porin-deficient clinical isolates decreased MICs of all carbapenems, but particularly of ertapenem, confirming that porin disruption disproportionately impairs ertapenem permeation.

The second concerns biosample SAMN05170257, an *Escherichia coli* isolate susceptible to tetracycline despite harbouring resistance to 13 other antibiotics. To explain the susceptibility, the teacher first used UniProt and GO to list the known ways bacteria resist tetracycline, which were then checked against the antibiogram. The broad-spectrum AcrAB efflux pump was ruled out because tigecycline, which AcrAB also exports, remained effective. Tetracycline-specific efflux by TetA and TetB was ruled out for the same reason, since these pumps spare tigecycline by design. With every known mechanism excluded, the teacher concluded that the isolate carries none of them and predicted susceptibility. This is consistent with Tuckman et al. [16], who attribute 93% of tetracycline resistance in *E. coli* clinical isolates to identifiable resistance genes: when none are present, susceptibility is the expected outcome. The teacher also noticed a clue the original investigators did not discuss. The biosample is resistant to gentamicin and tobramycin but susceptible to amikacin, a pattern consistent with intact outer-membrane porins and a substrate-restricted AAC(3) acetyltransferase. This means tetracycline can enter the cell freely and reach the ribosome—reinforcing the prediction of susceptibility through an argument the source study did not make.

### Ablation Study

The primary evaluation criterion for an LLM-based predictor such as BacteReason is the error rate (accuracy). Beyond this, we investigate whether teacher-generated rationales drive prediction improvement. An accuracy gain in the student LLM would constitute indirect evidence that the rationales encode mechanistically relevant information for susceptibility inference.

We compiled a benchmark of 200 question–answer pairs disjoint from the training data (see Methods) and compare BacteReason’s error rate against three ablated configurations:

1. BacteReason_w/o-Finetuning_: the base open-weight LLM with no fine-tuning.
2. BacteReason_w/o-Reasoning_: the same base LLM fine-tuned on question–answer pairs only, with rationales removed from the training data.
3. BacteReason_w/o-Database_: the same base LLM fine-tuned on full question–rationale– answer triplets, but with rationales generated by a teacher LLM that lacks TogoMCP access.

Figure 3(a) compares predicted and ground-truth susceptibility labels extracted from the final sentence of each model’s inference. See Methods for details about inference generation. The four models form a monotonic series in which each successive component lowers the error rate. The untuned baseline, BacteReason_w/o-Finetuning_, achieves an error rate of 32%. Comparing BacteReason_w/o-Reasoning_ (29%) against BacteReason_w/o-Finetuning_ (32%) isolates the contribution of fine-tuning on labels alone, a configuration functionally equivalent to training a conventional supervised classifier on the question–answer mapping. Comparing BacteReason_w/o-Database_ (24%) against BacteReason_w/o-Reasoning_ (29%) isolates the contribution of teacher-generated rationales, showing that mechanistic reasoning supervision contributes more than label supervision alone. Comparing BacteReason (18%) against BacteReason_w/o-Database_ (24%) isolates the contribution of knowledge-graph grounding, showing that rationales retrieved through TogoMCP outperform rationales generated without database access. In summary, these ablation studies demonstrate that both the presence of rationale supervision and the quality of the rationales are critical to prediction accuracy.

**Figure 3:**
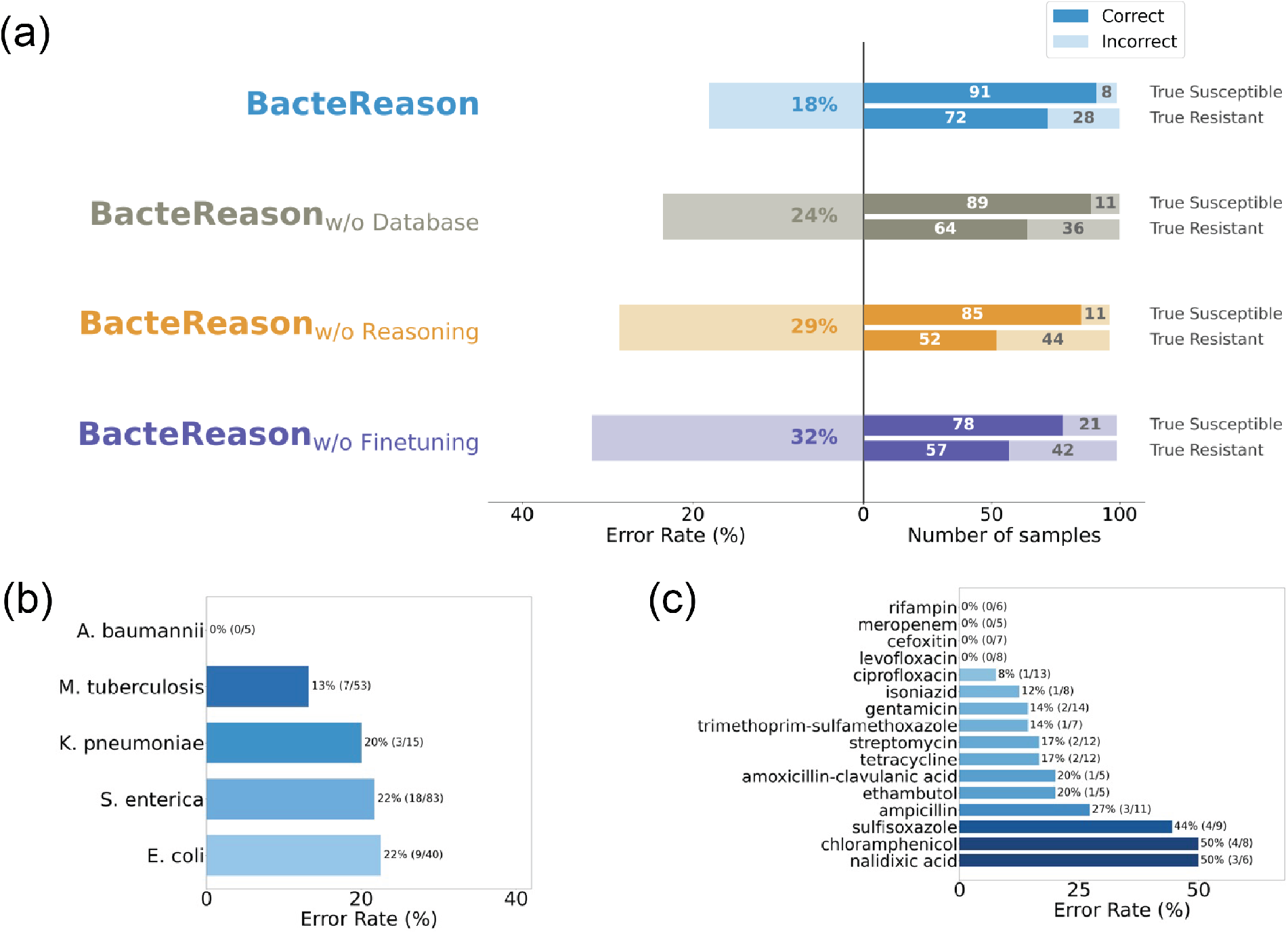
(a) Error rate comparison across four configurations of the same base LLM. BacteReason: open-weight LLM fine-tuned with the question–rationale–answer triplets. BacteReason_w/o-Finetuning_: untuned open-weight LLM. BacteReason_w/o-Reasoning_: the same open-weight LLM fine-tuned on question–answer pair only. BacteReason_w/o-Database_: the same base LLM fine-tuned on question–rationale–answer triplets whose rationales were generated by the teacher LLM without TogoMCP access. (b) Per-species and (c) per-antibiotic error rates for BacteReason.

To interpret BacteReason’s prediction pattern, we present its per-species and per-antibiotic error rates in Figure 3(b) and (c), respectively. For BacteReason, *E. coli, S. enterica* and *K. pneumoniae* are harder than *M. tuberculosis* and *A. baumannii*. Per antibiotic, BacteReason excels on *β*-lactams, and struggles on sulfonamides, chloramphenicol, and nalidixic acid.

### Topology Analysis

To inspect why rationales improve prediction, we apply the hidden-state topology analysis of Minegishi et al. [17] to three reasoning variants (BacteReason, BacteReason_w/o-Database_, and BacteReason_w/o-Finetuning_) during inference. We project their hidden-state representations into a 3D embedding and measure two topological properties of the resulting reasoning trajectory: the number of unique nodes visited and the graph diameter (i.e, longest shortest path).

Figure 4(a) shows one representative reasoning trajectory for each model for biosample SAMN18473032. For BacteReason, three turning points (steps at which the cluster assignment shifts most sharply) stand out: Step 1 (initial resistance-mechanism query), Step 7 (genotype information retrieval), and Step 37 (ontology-grounded *erm* gene annotation). All three coincide with TogoMCP interactions, hinting at why BacteReason ranges more widely than BacteReason_w/o-Database_.

**Figure 4:**
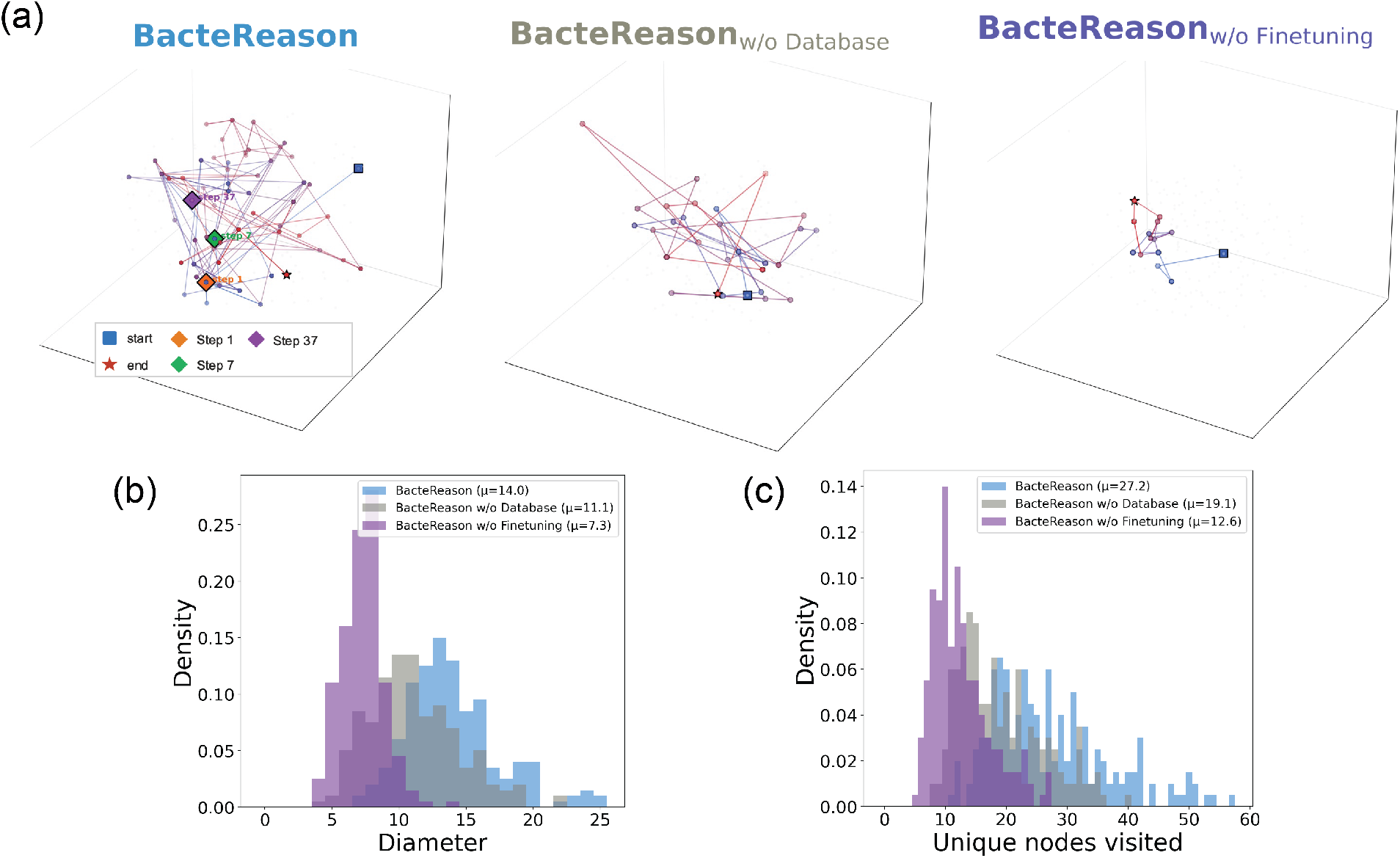
Topology analysis of the hidden states of three reasoning models during inference: BacteReason, BacteReason_w/o-Database_, and BacteReason_w/o-Finetuning_. (a) Representative reasoning-topology visualisation for biosample SAMN18473032. t-SNE projects the *K* = 200 shared centroids into 3D; node colour encodes temporal progression (blue → red). Coloured diamonds mark the top-3 turning points (steps with the largest cluster-centroid jump in shared PCA space). (b) Diameter (longest shortest path), capturing the spatial extent of reasoning. (c) Number of unique nodes visited, reflecting the breadth of the rationale.

Figure 4(b)–(c) show that this pattern holds across the benchmark. BacteReason visits more than twice as many unique nodes as BacteReason_w/o-Finetuning_ and traces nearly twice the graph diameter, suggesting that mechanistic rationales teach the model to canvas a wider region of representation space before committing to a prediction. BacteReason_w/o-Database_ sits between the two, indicating that knowledge-graph grounding amplifies this effect beyond what rationale supervision alone provides. Across all three variants, both topological metrics correlate negatively with prediction error rate: models that reason more broadly predict more accurately. Together, these results explain why reasoning supervision generalises: it pushes the model toward broader mechanistic exploration rather than shallow pattern matching. Topology analysis details are reported in Methods.

## Discussion

We developed BacteReason, a fine-tuned reasoning LLM specialised in predicting antibiotic susceptibility. Each question presents a clinical biosample’s antibiogram, together with the name of the target antibiotic; the answer is the ground-truth susceptibility label of the target antibiotic; and the rationale is generated by a proprietary teacher LLM prompted to explain the molecular mechanism underlying that label. The teacher LLM is interfaced to TogoMCP, a biomedical knowledge-graph retrieval tool. Our benchmark shows that fine-tuning with rationales improves prediction accuracy: BacteReason achieves an error rate of 18%, a relative improvement of 43% over the untuned baseline and 38% over the same base model fine-tuned without rationales. The topology analysis provides an explanation for this improvement: models trained with rationales explore broader regions of hidden-state space, engaging in richer deliberation rather than shallow pattern matching. These results indicate that reasoning supervision is the primary driver of correct generalisation in AMR prediction.

The stronger performance of BacteReason over BacteReason_w/o-Database_ (18% vs 24%) reflects the quality of the rationales used for fine-tuning. Understanding AMR mechanisms requires integrating diverse datasets—protein function, enzymatic reactions, drug– target interactions, gene ontology—that are distributed across separate public repositories. TogoMCP [12] addresses this challenge by providing a unified knowledge-graph interface to RDFPortal [18], a collection of biomedical databases including UniProt, ChEMBL, Reactome, and others. This enables the teacher LLM to compose queries across multiple databases within a single reasoning chain, producing more factually accurate rationales than unconstrained generation. This quality difference propagates through fine-tuning to yield more accurate predictions.

A key limitation of this study is that BacteReason predicts susceptibility from existing antibiogram patterns rather than anticipating future resistance acquisition; the model does not account for the temporal dynamics by which clinical isolates gain or lose resistance determinants under selective pressure. Addressing this will require training data that captures resistance-acquisition events: paired isolates from the same patient or lineage sampled before and after antibiotic exposure. The rationale-generation framework presented in this paper is in principle compatible with this extension: a teacher LLM could be prompted to generate rationales for why a given exposure pattern would select for a given resistance determinant, yielding question–rationale–answer triplets that supervise temporal prediction rather than static classification.

Three further directions could reduce the error rate. First, the data scaling curve (Supplementary Figure S1 (b)) has not saturated, suggesting that expanding the rationale dataset would directly improve performance. Second, the current implementation relies solely on supervised fine-tuning; applying group relative policy optimisation (GRPO), a reinforcement learning algorithm that optimises a model against a reward signal without a separate critic, could yield further gains by rewarding correct answers. Third, augmenting the input with genomic features would address BacteReason’s weakness on single-determinant phenotypes—cases where susceptibility or resistance hinges on a single gene that the antibiogram does not expose—by giving the model direct access to determinants invisible to the phenotype alone.

Because BacteReason derives its answer from a mechanistic chain of reasoning, correct predictions tend to indicate correct reasoning: when the model predicts well, the underlying rationale is more likely to have captured a relevant biological mechanism. This stands in contrast to black-box classifiers, where correct outputs reveal nothing about whether the model has learned the right causal structure or merely exploited a spurious correlation. BacteReason also addresses the interpretability bottleneck that has limited clinical adoption of machine learning for AMR prediction. Each prediction is accompanied by a self-contained molecular rationale that clinicians can inspect and use to inform treatment decisions, and that researchers can treat as mechanistic hypotheses for experimental validation. As biomedical knowledge graphs expand, LLM–ontology integration through protocols such as TogoMCP could extend the range of scientific questions that language models address with verifiable evidence. These directions point toward more capable reasoning LLMs for AMR prediction and, more broadly, for any prediction task in which correct generalisation depends on underlying mechanisms.

## Methods

### Fine-tuning

Supervised fine-tuning uses LoRA [19] (rank 16, *α* = 128, dropout 0.05) applied to all attention and feed-forward projections (q proj, k proj, v proj, o proj, gate proj, up proj, down proj). Training uses a cosine learning-rate schedule with peak rate 1 × 10^*−*4^, effective batch size 8 (micro-batch 1 × gradient accumulation 8), bfloat16 mixed precision with gradient checkpointing, and a maximum sequence length of 4,096 tokens for 10 epochs on a single NVIDIA A100 80 GB GPU, requiring approximately 15 seconds per training example. Hyperparameter sensitivity and data scaling analyses are provided in Supplementary Figure S1.

### Benchmark Data

Test biosamples are randomly selected from AMR Portal [11] subject to two constraints: (i) each biosample must have susceptibility labels for at least 10 antibiotics, and (ii) the heldout target labels comprise 100 susceptible and 100 resistant biosamples. Each question asks for the biosample’s susceptibility to a target antibiotic given its susceptibility labels against all remaining antibiotics. The answer is the ground-truth label of the held-out antibiotic: “susceptible” or “resistant.” The benchmark is constructed as an extrapolation benchmark: test biosamples are disjoint from those used for training, ensuring that no isolate appears in both splits and that predictions must generalise to previously unseen clinical isolates rather than being recovered from memorisation.

### Inference

Inference generation uses greedy decoding with a maximum output length of 16,384 tokens on a single NVIDIA A100 80 GB GPU, requiring approximately 5.7 minutes per example. The prediction is extracted from the last sentence of the output by keyword matching (“susceptible” or “resistant”); responses containing neither keyword within the token limit are excluded from accuracy computation.

### Topology Analysis

Following Minegishi et al. [17], we construct reasoning graphs by clustering hidden-state representations at each reasoning step. For each generated rationale, the text is split by newline into reasoning steps *r*_1_, *r*_2_, … , *r*_*T*_. A forward pass extracts hidden-state representations at layer 57 (90% depth of the 64-layer architecture), and each step’s representation is obtained by mean-pooling the token-level hidden states within that step. PCA reduces dimensionality from 5,120 to 50, and Mini-Batch *K*-means (*K* = 200) clusters the pooled representations. Each cluster centroid defines a node in the reasoning graph; consecutive assignments are connected as directed edges, yielding a per-sample graph. To enable direct comparison, all hidden-state representations from BacteReason, BacteReason_w/o-Database_, and BacteReason_w/o-Finetuning_ are pooled into a single shared clustering space before applying *K*-means, so that all three models trace paths through the same set of 200 nodes.

For visualisation, t-SNE (*n*_components_ = 3, perplexity= min(30, *K* ^−^ 1)) projects the *K* = 200 shared centroids into three dimensions. Each sample’s reasoning path is rendered as a directed trajectory, with edges coloured along a blue-to-red gradient indicating temporal progression. To identify salient reasoning transitions, we extract *turning points*: the top-3 steps at which the cluster assignment shifts to the most distant centroid. Formally, for a cluster sequence *c*_1_, *c*_2_, … , *c*_*T*_ , the jump distance at step *i* is 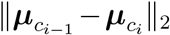, where ***µ***_*k*_ denotes the *k*-th centroid. Steps are ranked by this distance and the top-3 are designated turning points.

We report two graph-theoretic summary statistics per sample. *Diameter* is the longest shortest path between any two visited nodes, capturing the spatial extent of reasoning traversal. *Unique nodes* counts the number of distinct clusters visited, reflecting the breadth of the rationale. Together, these metrics quantify whether a model engages in broad, far-ranging reasoning or collapses to a repetitive, locally confined trajectory.

## Code and Data Availability

The fine-tuned BacteReason model weights are available on Hugging Face at https://huggingface.co/Playingyoyo/BacteReason. An interactive demo is hosted as a Hugging Face Space (https://huggingface.co/spaces/Playingyoyo/BacteReason). Code for reproducing the results is available as a Google Colab notebook at https://colab.research.google.com/drive/1BEwfxVyz3L3TAv1xObuC5Ef8oPH5oUGn?usp=sharing.

## Acknowledgements

This study is supported by JST MIRAI JPMJMI24H2.

## Author Contributions

All authors conceived the original idea. Y.O., A.R.K., and S.K. prepared the dataset. Y.D. reviewed the dataset. Y.O., R.T., and K.T. wrote the original manuscript. All authors discussed the results, commented on the manuscript, and approved the final version.

## References

[1] Murray, C. J. L., Ikuta, K. S., Sharara, F. et al. Global burden of bacterial antimicrobial resistance in 2019: a systematic analysis. Lancet 399, 629–655 (2022).

[2] Teillant, A., Gandra, S., Barter, D., Morgan, D. J. & Laxminarayan, R. Potential burden of antibiotic resistance on surgery and cancer chemotherapy antibiotic prophylaxis in the USA: a literature review and modelling study. The Lancet Infectious Diseases 15, 1429–1437 (2015).

[3] Tabak, Y. P. et al. Blood Culture Turnaround Time in U.S. Acute Care Hospitals and Implications for Laboratory Process Optimization. Journal of Clinical Microbiology 56, e00500–18 (2018).

[4] Yelin, I. et al. Personal clinical history predicts antibiotic resistance of urinary tract infections. Nature Medicine 25, 1143–1152 (2019).

[5] Kanjilal, S. et al. A decision algorithm to promote outpatient antimicrobial stewardship for uncomplicated urinary tract infection. Science Translational Medicine 12, eaay5067 (2020).

[6] Wei, J. et al. Chain-of-Thought Prompting Elicits Reasoning in Large Language Models. arXiv preprint 2201.11903 (2023).

[7] Kang, M., Lee, S., Baek, J., Kawaguchi, K. & Hwang, S. J. Knowledge-Augmented Reasoning Distillation for Small Language Models in Knowledge-Intensive Tasks. arXiv preprint 2305.18395 (2023).

[8] Kwon, T. et al. Large Language Models are Clinical Reasoners: Reasoning-Aware Diagnosis Framework with Prompt-Generated Rationales. arXiv preprint 2312.07399 (2024).

[9] Fallahpour, A. et al. BioReason: Incentivizing Multimodal Biological Reasoning within a DNA-LLM Model. arXiv preprint 2505.23579 (2025).

[10] Ho, N., Schmid, L. & Yun, S.-Y. Large Language Models Are Reasoning Teachers. arXiv preprint 2212.10071 (2023).

[11] Jesudason, T. EMBL–EBI’s AMR portal: a new gateway in global antimicrobial resistance research. The Lancet Microbe 7 (2026).

[12] Kinjo, A. R., Yamamoto, Y., Bustamante-Larriet, S., Labra-Gayo, J.-E. & Fujisawa, T. TogoMCP: Natural Language Querying of Life-Science Knowledge Graphs via Schema-Guided LLMs and the Model Context Protocol. bioRxiv preprint 2026.03.19.713030 (2026).

[13] Anthropic. Claude Opus 4.5 (2025). URL https://www.anthropic.com.

[14] Qwen Team. QwQ-32B: Embracing the Power of Reinforcement Learning (2025). URL https://qwenlm.github.io/blog/qwq-32b/.

[15] Doumith, M., Ellington, M. J., Livermore, D. M. & Woodford, N. Molecular mechanisms disrupting porin expression in ertapenem-resistant Klebsiella and Enterobacter spp. clinical isolates from the UK. The Journal of Antimicrobial Chemotherapy 63, 659–667 (2009).

[16] Tuckman, M. et al. Occurrence of Tetracycline Resistance Genes among Escherichia coli Isolates from the Phase 3 Clinical Trials for Tigecycline. Antimicrobial Agents and Chemotherapy 51, 3205–3211 (2007).

[17] Minegishi, G., Furuta, H., Kojima, T., Iwasawa, Y. & Matsuo, Y. Topology of Reasoning: Understanding Large Reasoning Models through Reasoning Graph Properties. arXiv preprint 2506.05744 (2025).

[18] Kawashima, S., Katayama, T., Hatanaka, H., Kushida, T. & Takagi, T. NBDC RDF portal: a comprehensive repository for semantic data in life sciences. Database: The Journal of Biological Databases and Curation 2018, bay123 (2018).

[19] Hu, E. J. et al. LoRA: Low-Rank Adaptation of Large Language Models. arXiv preprint 2106.09685 (2021).

