## Supplementary information for "BacteReason: A Reasoning Model for Antimicrobial Resistance Prediction"

### Supplementary Text S1: Example Prompt

You are given a Escherichia coli clinical isolate (BioSample SAMN16435623) tested against multiple antibiotics. Phenotype profile: susceptible to chloramphenicol ($= 8$ mg/L), trimethoprim-sulfamethoxazole ($\leq 0.12$ mg/L), sulfisoxazole ($\leq 16$ mg/L), cefoxitin ($= 4$ mg/L), gentamicin ($= 0.5$ mg/L), kanamycin ($\leq 8$ mg/L), nalidixic acid ($= 2$ mg/L), streptomycin ($\leq 32$ mg/L), tetracycline ($\leq 4$ mg/L), ceftiofur ($= 0.5$ mg/L), ampicillin ($= 2$ mg/L), amoxicillin-clavulanic acid ($= 4$ mg/L), ceftriaxone ($\leq 0.25$ mg/L) and azithromycin ($= 2$ mg/L).

**Instruction used during reasoning traces construction**

Develop a detailed, evidence-based explanation for the molecular mechanism(s) most likely responsible for the biosample's susceptibility to ciprofloxacin, using the biosample's responses to other antibiotics as diagnostic clues. Proceed as follows: (1) Profile-driven inference: Identify phenotypic patterns across antibiotic classes. (2) Iterative hypothesis testing: Generate hypotheses, query RDFPortal databases (PubChem, ChEBI, ChEMBL, PDB, UniProt, GO, Reactome, Rhea), state whether evidence supports or contradicts each hypothesis, and repeat until one survives. (3) Final mechanism presentation: Present the surviving hypothesis as step-by-step reasoning tied to phenotype patterns and verified molecular evidence. Conclude exactly with: "Therefore, BioSample SAMN16435623 is susceptible to ciprofloxacin."

⇓ *Replaced during fine-tuning and inference* ⇓

**Instruction used during fine-tuning and inference**

Predict whether BioSample SAMN16435623 is susceptible or resistant to ciprofloxacin. Conclude exactly with: "Therefore, BioSample SAMN16435623 is susceptible/resistant to ciprofloxacin."

**Supplementary Text S2: BacteReason's Response to SAMEA104075919's Susceptibility to Meropenem**

### Hypothesis 1: *bla*$_{\text{NDM-5}}$ is silenced or minimally expressed

**Rationale.** The carbapenemase gene may be present but transcriptionally inactive due to promoter disruption, insertion-sequence interference, or unfavorable genetic context.

**Evidence.** AMRPortal HMM-based annotation detects the complete ORF with high-confidence evidence linked to NCBI Pathogen Detection. The gene was detected in multiple copies, consistent with active mobile-element carriage.

**Verdict.** *Contradicted.* HMM detection requires substantial coding-region conservation; truncation would fail recognition. More importantly, ertapenem resistance requires active NDM-5 enzyme, proving expression.

### Hypothesis 2: NDM-5 has intrinsically reduced catalytic efficiency against meropenem

**Rationale.** Structural differences between carbapenems could produce differential substrate binding and hydrolysis rates.

**Evidence.** ChEBI shows ertapenem (CHEBI:404903; $C_{22}H_{25}N_3O_7S$; MW $\approx$ 475.5) carries a bulky 3-carboxyphenyl substituent; meropenem (CHEBI:43968; $C_{17}H_{25}N_3O_5S$; MW $\approx$ 383.5) has a smaller dimethylcarbamoyl-pyrrolidinyl side chain. UniProt C7C422 establishes NDM hydrolysis of penicillins, cephalosporins, and carbapenems. NDM-5 differs from NDM-1 by V88L and M154L substitutions, which enhance carbapenem hydrolysis.

**Verdict.** *Contradicted.* NDM-5 is hyperactive relative to NDM-1; the differential phenotype cannot be explained by intrinsic specificity.

### Hypothesis 3: Outer-membrane porin modifications create differential permeability barriers

**Rationale.** Carbapenems must traverse OmpK35/OmpK36 to reach periplasmic $\beta$-lactamases; porin alterations could differentially restrict entry based on molecular size and charge.

**Evidence.** UniProt confirms OmpK35/OmpK36 as primary porins in *K. pneumoniae* (variants S4SBZ2, S4SBW9, A0A2P1E303). GO:0015288 defines porin transfer as $< 1000\,\mathrm{Da}$ via $\beta$-barrel channels; ertapenem's larger size and 3-carboxyphenyl charge profile reduce permeation through narrowed channels. NCBI Gene records OmpK36 variants known to cause selective carbapenem resistance.

**Verdict.** *Supported, but incomplete.* Porin modifications explain differential entry but cannot alone overcome active NDM-5 periplasmically.

### Hypothesis 4: Pharmacokinetic differences compound the porin-permeability effect

**Rationale.** Ertapenem exhibits $\sim$95% serum protein binding versus $\sim$2% for meropenem, producing dramatic differences in free drug concentration available for periplasmic uptake.

**Evidence.** ChEBI shows ertapenem's ionizable carboxylate increases albumin affinity; meropenem's dimethylcarbamoyl lacks this functionality. Rhea (RHEA:18693) and GO:0008800 establish $\beta$-lactamase activity as concentration-dependent, with substrate availability rate-limiting.

**Verdict.** *Supports the integrated model.*

#### Hypothesis 5 (Final): Dual-barrier model with pharmacokinetic amplification

**Integrated rationale.** NDM-5 is expressed and functional, as proven by ertapenem resistance and cefoxitin hydrolysis. Porin modifications (likely OmpK36) reduce carbapenem entry, affecting the larger ertapenem more than meropenem. Protein-binding differences reduce free ertapenem to ∼5% versus ∼98% for meropenem. Combined, periplasmic meropenem exceeds NDM-5 hydrolytic capacity and reaches PBPs at bactericidal concentrations; ertapenem does not.

**Supporting cross-database evidence.** UniProt OmpK36 variants confirm that porin mutations accumulate under antibiotic pressure in *K. pneumoniae*. ChEBI confirms the ∼25% MW difference and distinct electrostatic profiles. GO annotations establish concentration-dependence of both porin-mediated transport (GO:0015288) and $\beta$-lactamase activity (GO:0008800). Aminoglycoside susceptibility (amikacin, gentamicin, tobramycin) confirms the outer membrane is not globally impermeable. Tigecycline susceptibility despite *oqxAB* indicates efflux is not the primary determinant.

### Final Mechanism Inference

The molecular basis for SAMEA104075919's susceptibility to meropenem despite carrying $bla_{\text{NDM-5}}$ is a **pharmacokinetic–structural dual-barrier effect**.

**Step 1.** NDM-5 carbapenemase is actively expressed. Ertapenem and cefoxitin resistance confirm enzyme function; only NDM-5 explains the cephamycin phenotype, because CTX-M-14 lacks cephamycinase activity and SHV has minimal cephamycin activity.

**Step 2.** Outer-membrane porin alterations restrict carbapenem entry. OmpK36 mutations or reduced OmpK35/OmpK36 expression create a size-selective barrier favoring meropenem (383 Da, compact pyrrolidinyl) over ertapenem (476 Da, bulky carboxyphenyl).

**Step 3.** Differential protein binding amplifies the permeability effect. With ∼95% versus ∼2% binding, ∼20-fold more free drug is available for meropenem before considering permeability.

**Step 4.** Meropenem achieves bactericidal periplasmic concentrations. High free-drug fraction and better porin penetration allow accumulation faster than NDM-5 can hydrolyze it, saturating enzyme turnover ($k_{\text{cat}}$) and reaching PBPs.

**Step 5.** Ertapenem fails to reach threshold. The multiplicative protein-binding × porin-entry deficit holds periplasmic concentrations below the level needed to saturate NDM-5; the drug is hydrolyzed before acting.

**Why alternative hypotheses failed.** Gene silencing is refuted by ertapenem resistance proving enzyme activity. The enzyme-specificity hypothesis is refuted because NDM-5 has enhanced, not reduced, carbapenem hydrolysis. Simple porin loss is insufficient because it cannot explain preservation of any carbapenem against an active MBL.

**Acceptance rationale.** The dual-barrier model uniquely explains (i) resistance to ertapenem but not meropenem despite identical enzymatic target, (ii) pan-$\beta$-lactam resistance consistent with active NDM-5, (iii) aminoglycoside susceptibility confirming membrane integrity, and (iv) the correlation between molecular properties (size, protein binding) and phenotype. The model integrates evidence from AMRPortal (phenotype/genotype), ChEBI (molecular structures), UniProt (porin biology, enzyme function), and Gene Ontology (mechanistic definitions).

---

Therefore, SAMEA104075919 is susceptible to meropenem.

**Supplementary Text S3: BacteReason's Response to SAMN05170257's Susceptibility to Tetracycline**

### Hypothesis 1: AcrAB-TolC efflux pump upregulation

**Rationale.** RND-family AcrAB-TolC exports tetracyclines, fluoroquinolones, and selected $\beta$-lactams. Given fluoroquinolone resistance, upregulated efflux is plausible.

**Evidence.** UniProt P31224, P0AE06, P02930 confirm AcrAB-TolC substrate range including tetracyclines, macrolides, and fluoroquinolones, but limited aminoglycoside export due to polycationic character. Tigecycline—designed to evade TetA/TetB—remains a partial AcrAB substrate.

**Verdict.** *Contradicted.* Upregulated AcrAB-TolC would reduce susceptibility to both tetracycline and tigecycline, contradicting the observed double-susceptible phenotype. The selective aminoglycoside pattern also requires an enzymatic, not efflux, mechanism.

### Hypothesis 2: TetA/TetB present but defective

**Rationale.** MFS-family TetA/TetB efflux pumps are tetracycline-specific (excluded by tigecycline's 9-*tert*-butylglycylamido group). A non-functional copy could produce a susceptible phenotype.

**Evidence.** UniProt A0A037Y375, B1VCE9 describe TetA/TetB as proton-antiport transporters; Rhea RHEA:35235 gives the efflux reaction. PubChem CID 54686904 confirms tigecycline's steric exclusion from TetA/TetB.

**Verdict.** *Contradicted.* Even partially functional TetA/TetB would yield tetracycline resistance with tigecycline susceptibility, not the observed double-susceptible phenotype. Gene absence, not gene dysfunction, is the simpler explanation.

### Hypothesis 3: Ribosomal protection proteins (TetM /TetO /TetW)

**Rationale.** GTPase-mediated ribosomal protection displaces tetracycline from the 30S A-site, conferring resistance to classical tetracyclines and reduced tigecycline susceptibility.

**Evidence.** UniProt Q02652 (TetM), P10952 (TetO); PDB 3J9Y shows TetM bound to the *E. coli* ribosome; PDB 4V6V shows TetO. These mechanisms invariably confer tetracycline resistance.

**Verdict.** *Contradicted.* Tetracycline susceptibility excludes operative ribosomal protection.

### Hypothesis 4: Tetracycline-inactivating enzymes (TetX-type monooxygenases)

**Rationale.** TetX catalyzes NADPH-dependent C11a hydroxylation, destroying all tetracyclines including tigecycline.

**Evidence.** Rhea RHEA:50004 documents the reaction: tetracycline + NADPH + $O_2$ + $H^+$ $\rightarrow$ 11a-hydroxytetracycline + $NADP^+$ + $H_2O$. TetX is uncommon in *E. coli* (chiefly *Bacteroides*).

**Verdict.** *Contradicted.* Enzymatic destruction would yield resistance, not susceptibility.

### Hypothesis 5 (Final): Absence of tetracycline-resistance determinants

**Rationale.** The concordant susceptibility to tetracycline and tigecycline, combined with mechanism-specific resistances for other drug classes, indicates that this isolate lacks any tetracycline-targeting resistance determinant.

**Cross-database evidence.** The ESBL pattern is consistent with UniProt P28585 and A0A5R8T042, which confirm CTX-M hydrolysis of oxyimino-cephalosporins but not cephamycins or carbapenems—matching the observed $\beta$-lactam phenotype. The AAC(3) signature is corroborated by PDB 7MQM, 6MN5, 9MH6, which show acetylation of the 3-amino group of gentamicin/tobramycin, while amikacin's 1-$N$-HABA side chain sterically blocks enzyme access—explaining the selective aminoglycoside pattern without invoking permeability loss. The Qnr/GyrA signature is supported by UniProt A0A1D8ELK6, which describes QnrS protection of DNA gyrase, a mechanism isolated to quinolones. For plasmid epidemiology, in *E. coli* the genes $bla_{\text{CTX-M}}$, *aac(3)*, *qnrS*, and *sul1*/*sul2* commonly co-locate on IncF plasmids; *tet* determinants are not obligate cargo and their presence is variable.

**Verdict.** *Supported.* All non-tetracycline resistances have specific, target-restricted mechanisms; none extends to tetracyclines. The susceptibility to both tetracycline and tigecycline indicates no *tet* determinant of any class.

### Final Mechanism Inference

**Step 1: Membrane permeability is intact.** Differential aminoglycoside susceptibility (amikacin S; gentamicin, tobramycin R) requires intact porin uptake and inner-membrane energization. OmpF/OmpC loss or proton-motive-force defects would suppress all aminoglycosides. Tetracyclines enter *E. coli* via OmpF/OmpC; intact permeability predicts unimpeded access to the ribosomal target.

**Step 2: Enzymatic capacity is substrate-restricted.** The AAC(3) signature acetylates 3-amino groups of gentamicin/tobramycin, excluded from amikacin by its HABA side chain. *E. coli* has no known acetyltransferase or analogous enzyme that modifies the tetracycline scaffold.

**Step 3: Efflux is not operative for tetracyclines.** Fluoroquinolone resistance is explained by GyrA/ParC mutations or Qnr rather than AcrAB-TolC overexpression: tigecycline (an AcrAB substrate) remains fully susceptible, ruling out major RND upregulation. Tetracycline-specific TetA/TetB are excluded by tigecycline susceptibility (absence of class-specific efflux).

**Step 4: 30S ribosomal target is unmodified.** Azithromycin resistance reflects 50S-targeting mechanisms (*mef* efflux or *erm* methylation), mechanistically isolated from the 30S A-site where tetracyclines bind. Absence of *tetM/tetO/tetW* is indicated by the phenotype, since ribosomal protection invariably produces tetracycline resistance.

**Step 5: Mechanistic integration.** The resistome is a coherent constellation typical of clinical MDR *E. coli*—ESBL (∼CTX-M), AAC(3), GyrA/ParC or Qnr, Dfr variant, and likely macrolide efflux. These genes commonly co-segregate on IncF plasmids but not obligately with *tet* determinants. The absence of *tetA/tetB/tetC/tetD* (efflux), *tetM/tetO/tetW* (ribosomal protection), and *tetX* (enzymatic degradation) means there is no operative tetracycline-resistance pathway.

**Step 6: Conclusion.** Tetracycline enters via intact OmpF/OmpC porins, accumulates without efflux-mediated export, and binds the 30S A-site without displacement or enzymatic destruction. It blocks aminoacyl-tRNA accommodation and exerts bacteriostatic activity.

**Why alternative hypotheses failed.** AcrAB-TolC upregulation is refuted by tigecycline susceptibility (AcrAB substrate). Defective TetA/TetB is refuted by tigecycline susceptibility (expected with TetA/TetB) and by parsimony favoring gene absence. Ribosomal protection is refuted because TetM/TetO/TetW invariably confer tetracycline resistance. TetX inactivation is refuted because enzymatic destruction produces resistance, not susceptibility.

**Acceptance rationale.** The "absent determinant" model uniquely explains the full profile: (i) target-restricted mechanisms for each non-tetracycline resistance, (ii) preserved permeability evidenced by amikacin susceptibility, (iii) absence of class-wide efflux evidenced by tigecycline susceptibility, and (iv) the well-documented variability of *tet* cargo on IncF resistance plasmids. Evidence integrates AMRPortal (phenotype/genotype), UniProt (transporters and enzymes), PDB (structural mechanisms), Rhea (enzymatic reactions), and PubChem (small-molecule geometry).

---

Therefore, SAMN05170257 is susceptible to tetracycline.

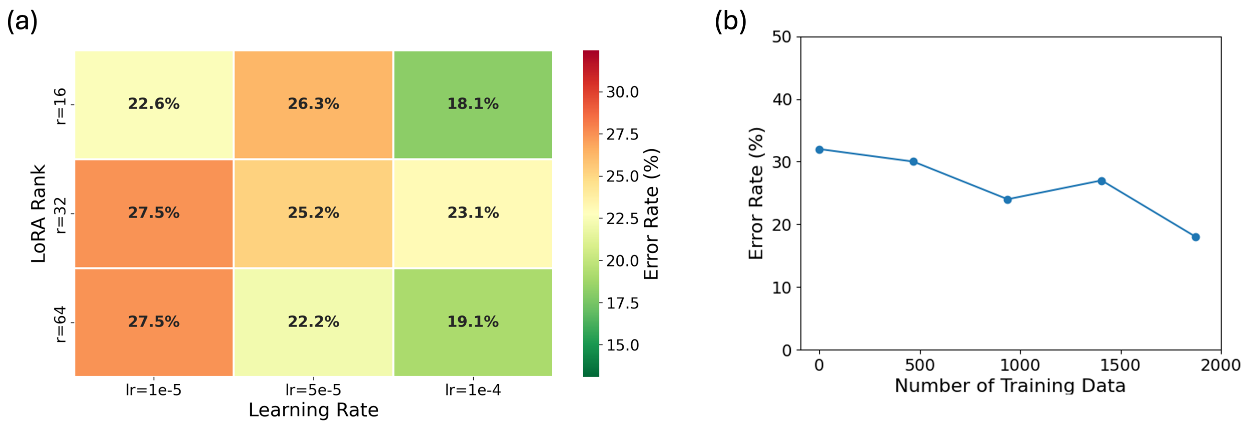


Supplementary Figure S1: Fine-tuning details: (a) LoRA hyperparameter sensitivity (error rate across rank and learning-rate configurations), and (b) effect of training data size on error rate, showing steady improvement with no clear saturation at 1,875 samples.
